# γδ T cell-mediated cytotoxicity against patient-derived healthy and cancer cervical organoids

**DOI:** 10.1101/2023.07.25.550464

**Authors:** Junxue Dong, David Holthaus, Christian Peters, Stefanie Koster, Marzieh Ehsani, Alvaro Quevedo-Olmos, Hilmar Berger, Michal Zarobkiewicz, Mandy Mangler, Rajendra Kumar Gurumurthy, Nina Hedemann, Cindrilla Chumduri, Dieter Kabelitz, Thomas F. Meyer

## Abstract

Cervical cancer is a leading cause of death among women globally, primarily driven by high-risk papillomaviruses. However, the effectiveness of chemotherapy is limited, underscoring the potential of personalized immunotherapies. Patient-derived organoids, which possess cellular heterogeneity, proper epithelial architecture and functionality, and long-term propagation capabilities offer a promising platform for developing viable strategies.

In addition to αβ T cells and natural killer (NK) cells, γδ T cells represent a cell population with significant therapeutic potential against both hematologic and solid tumours.

To evaluate the efficacy of γδ T cells in cervical cancer treatment, we generated patient-derived healthy and cancer ectocervical organoids. Furthermore, we examined transformed healthy organoids, expressing HPV16 oncogenes E6 and E7. We analysed the effector function of *in vitro* expanded γδ T cells upon co-culture with organoids. Our findings demonstrated that healthy cervical organoids were less susceptible to γδ T cell-mediated cytotoxicity compared to HPV-transformed organoids and cancerous organoids.

To identify the underlying pathways involved in this observed cytotoxicity, we performed bulk-RNA sequencing on the organoid lines, revealing differences in DNA-damage and cell cycle checkpoint pathways, as well as transcription of potential γδ T cell ligands. We validated these results using immunoblotting and flow cytometry. We also demonstrated the involvement of BTN3A1 and BTN2A1, crucial molecules for γδ T cell activation, as well as differential expression of PDL1/CD274 in cancer, E6/E7+ and healthy organoids. Interestingly, we observed a significant reduction in cytotoxicity upon blocking MSH2, a protein involved in DNA mismatch-repair.

In summary, we established a co-culture system of γδ T cells with cervical cancer organoids, providing a novel *in vitro* model to optimize innovative patient-specific immunotherapies for cervical cancer.

## Introduction

According to latest WHO data, cervical cancer is the fourth leading cause of death among women worldwide ^1^. The majority of cervical cancer are strongly associated with persistent infection with high-risk oncogenic human papilloma virus (HR-HPV) ^2,3^. Among HR-HPVs, HPV16 and HPV18 account for about 70% of cervical cancers ^4^. Yet, additional infectious agents have also been implicated as co-drivers of cervical cancer ^5^. Considering that only a quarter of cervical cancer patients respond to chemotherapy, the development of more personalized therapies is urgently needed. Recent advances in organoid technology have enabled the culture of human ectocervical organoids that faithfully replicate healthy and cancerous tissues ^6^. These organoids serve as a promising *in vitro* model, preserving original cellular heterogeneity and accurately recapitulating epithelial architecture and functionality. Recognizing the absence of *in vitro* models for HPV, we have earlier reported the transformation of healthy cervical organoids with HPV-derived oncogenes E6 and E7 *via* lentiviral transfer ^5^. These transformed organoids exhibit significant deviations from their corresponding healthy controls ^5^.

A numerically minor yet important subset of T lymphocytes in the peripheral blood endowed with anti-cancer activity is γδ T cells ^7,8^. γδ T cells differ from conventional αβ T cells in at least two important aspects: (i) γδ T cells do not recognize peptides presented by MHC class I or class II molecules but rather recognize phosphorylated intermediates (“phosphoantigens”) of the cholesterol synthesis pathway and additional unconventional ligands; and (ii) γδ T cells do not require MHC molecules for antigen recognition, which opens the possibility of applying γδ T cells across HLA barriers ^9,10^. γδ T cells express CD3-associated T cell receptors (TCR) composed of γ and δ chains ^11,12^. Like other cytotoxic effectors, γδ T cells directly participate in the elimination of tumour cells, but they also indirectly control the tumour immune response by modulating the activity and functions of other immune cells ^7^. In healthy adults, approximately 1-5% of circulating T lymphocytes express Vδ2 paired with Vγ9; the proportion of such Vδ2 T cells can rapidly increase during the acute phase of several infectious disease ^11,12^. Recent studies have shed light on the mode of recognition of phosphoantigens (pAg) by γδ T cells. In this context, an indispensable role of members of the butyrophilin (BTN) family of transmembrane molecules has been discovered. According to the current model, microbe- or tumour-derived pyrophosphates bind to the intracellular B30.2 domain of BTN3A1 which then interacts with BTN2A1 to induce a conformational alteration of the extracellular BTN3A1/2A1 complex which is recognized by the γδ TCR ^13–16^.

γδ T cells have well-established protective roles in cancer ^8,17,18^. Multiple mouse models have demonstrated their protective role in cancer progression ^18,19^. However, previous studies so far have only investigated the effect of γδ T cells on cervical cancer-derived cancer cell lines and tissue slices ^18,20^ or utilized NK cells ^21^, while studies comparing their effects on patient-derived organoids have not yet been reported. In this study, we provide a model to assess γδ T cell-mediated cytotoxicity by live cell imaging and flow cytometry-based assays.

## Results

### Ectocervical organoids accurately represent the phenotype of the parental tissue

To establish a suitable model mimicking ectocervical tissue, we have established patient-derived organoids (PDOs) cervical tissues in our earlier studies ^5,6,22^. These lines were grown in T25 tissue flasks on irradiated 3T3-J2 mouse fibroblast feeder cells and su8bsequently embedded in Matrigel to generate 3D organoids (Fig. 1A). In order to create paired HPV E6E7-positive lines (referred to as HPV+), the healthy organoid lines were transformed with a construct containing human papilloma virus (HPV) oncogenes E6 and E7 as described earlier ^5,22^. We were able to passage all lines for approximately 20 passages prior to growth arrest. Visual inspection under brightfield microscopy revealed no discernible phenotypic difference between healthy, HPV+ and cancer lines (Fig. 1B). Amplification of integrated HPV16 E6E7 oncogenes was detected by PCR of genomic DNA in the HPV+ and cancer organoids (Fig. 1C). Since HPV16 and 18 are the most common causes of cervical cancers, we also assessed the HPV18 status in our organoids. HPV18 was not detected in any of the lines; primer sensitivity was verified using HeLa cells as a positive control (Supplementary Fig. 1).

**Figure 1:**
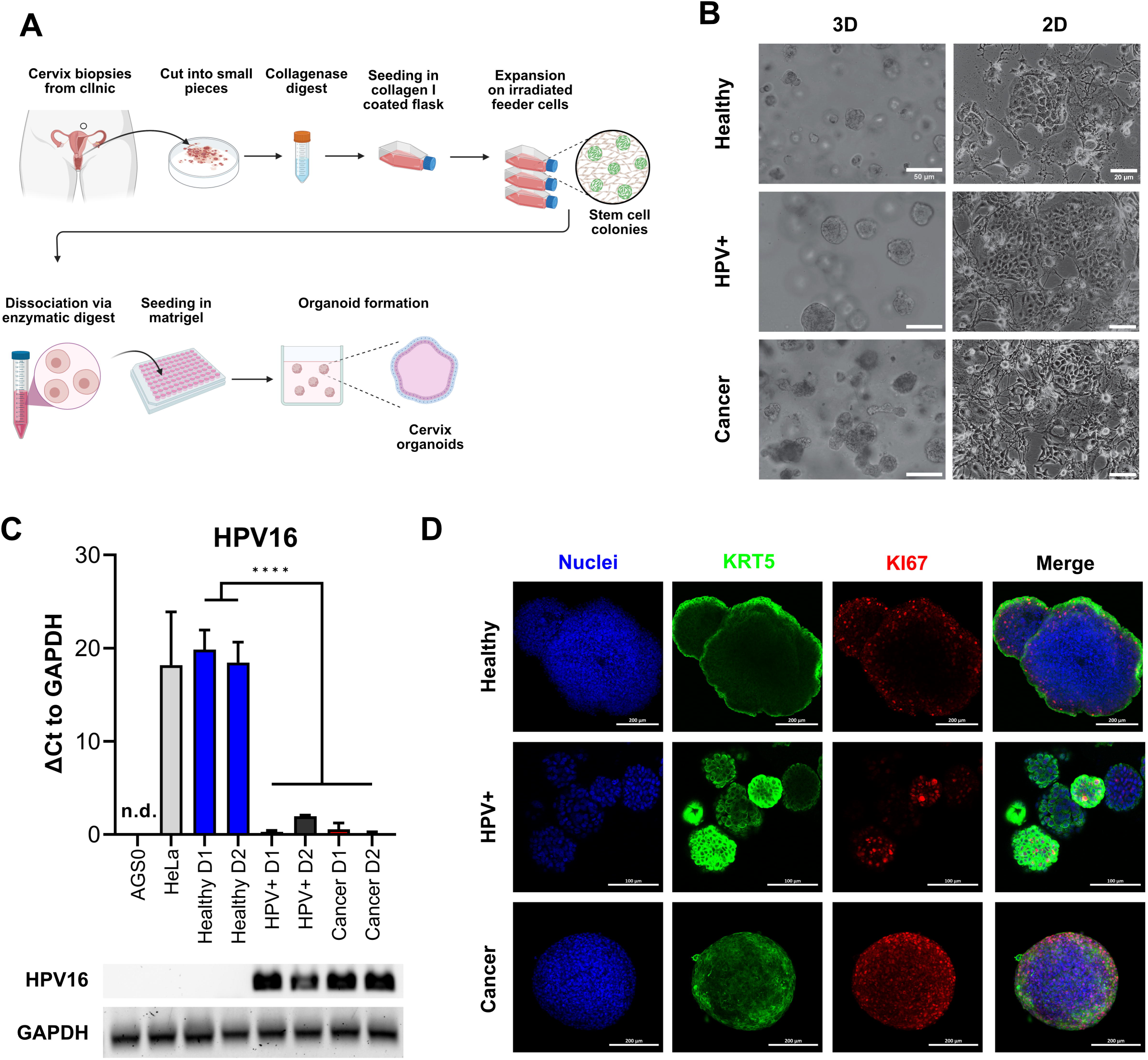
Characterization of healthy and cancer-derived ectocervical organoid cultures. A) Schematic representation of the establishment of organoids derived from ectocervical tissue. B) Representative brightfield images of three-dimensional (3D) and two-dimensional (2D) cultures of healthy ectocervical organoids, HPV oncogene E6E7-transformed paired organoids (HPV+) and cancer-derived organoids. Scale bars represent 50 µm for 3D organoids and 20 µm for 2D cultures. C) HPV16 status of the patient-derived organoid cultures. Genomic DNA was amplified by RT-qPCR and ΔCt values were calculated with GAPDH as a loading control. Agarose gel electrophoresis was performed to determine HPV status. AGS0 and HeLa cells served as negative controls. HPV+ and cancer organoids showed the presence of integrated HPV oncogenes E6E7. RT-qPCR experiments show mean (± 95% CI) from ≥ 3 technical replicas of two independent biological replicates. n.d., not detectable. Statistical significance between conditions was determined using a Two-Way ANOVA with Tukey’s correction for multiple testing. **** p < 0.0001 D) Representative immunofluorescent images of 3D organoids. Nuclei were stained with Hoechst33342 (blue), ectocervical tissue with cytokeratin 5 (KRT5, green) and proliferating cells with marker of proliferation 67 (KI67, red). Scale bars represent 100 µm for HPV+ organoids and 200 µm for healthy and cancer organoids.

To validate the ectocervical nature of our organoids, we next characterized the organoid lines for marker gene expression and localization by whole mount immunofluorescence assays. All lines were positive for cytokeratin 5 (KRT5), a marker of ectocervix, in the outer layers of the organoids resembling the *in vivo* stratified squamous epithelium of the parental tissue. As the basal layer is responsible for proliferation *in vivo*, marker of proliferation KI67 (KI67)-positive cells are expected in this layer. Interestingly, KI67^+^ cells were indeed detected in the parabasal layers of healthy PDOs, while they were located throughout the whole organoid for HPV+ and cancer-derived lines (Fig. 1D). A more detailed characterization of ectocervical organoid cultures has been previously provided in our studies ^5,6^. Thus, the organoids reflect the *in vivo* phenotypes of healthy and cancerous ectocervical tissues.

### γδ T cells exhibit enhanced killing of HPV+ and cancer-derived organoids

γδ T cells are an important subset of T lymphocytes involved in anti-tumour immunity. They can kill tumour target cells by the secretory pathway *via* release of perforin and granzymes as well as by death receptor pathways (e.g. Fas/Fas-ligand) ^11^. To first assess whether the tissue identity is also reflected in the immune response of tumour-killing T cell subsets, we co-incubated *in vitro* expanded γδ T cells with healthy, HPV+ and cancer-derived organoid lines. Before co-incubation, the purity of γδ T-cell lines was assessed; only cultures with > 90% γδ T cells were used. Representative examples of purity determinations as analysed by flow cytometry are presented in Supplementary Fig. 2. To determine γδ T cell-mediated cytotoxicity, organoids were stained with Nucred Dead 647 (Yellow: general stain, Red: dead cells) as previously described for breast cancer-PDOs ^23^. This staining enabled the detection of an increasing number of dead cells within the organoid population by automatized live cell imaging. Live cell imaging revealed distinct responses of γδ T cells to HPV+ and cancer-derived organoid lines (Fig. 2A-D). The average accumulation of red stained dead cells in these groups was higher in comparison with healthy organoids (Fig. 2A, B). We observed a significant increase in the ratio of red/dead to yellow/live signal in cancer and HPV+ organoids in comparison to healthy organoids after 4 hours after analysing single organoids (Fig. 2C). When plotting the data as scatter plots a clear shift to the red/dead signal was observable in cancer and HPV+ organoids (Fig. 2D). Consistently, additional activation of γδ T cells by the pAg BrHPP resulted in enhanced cytotoxicity in all organoid lines (Supplementary Fig. 3). Scatters plots for untreated, unstained and dead cells are provided in Supplementary Fig. 3. To further confirm the findings, organoid-derived cells were co-incubated with γδ T cells and cytotoxicity was assessed by an alternative, protease-based cytotoxicity assays after 4h. Higher cytotoxicity was again observed in cancer PDOs after co-incubation with unstimulated γδ T cells (Fig. 2E, left), while treatment with BrHPP additionally resulted in a significant increase in γδ T cell-mediated cytotoxicity against HPV-transformed organoids and cancer lines (Fig. 2E, right).

**Figure 2:**
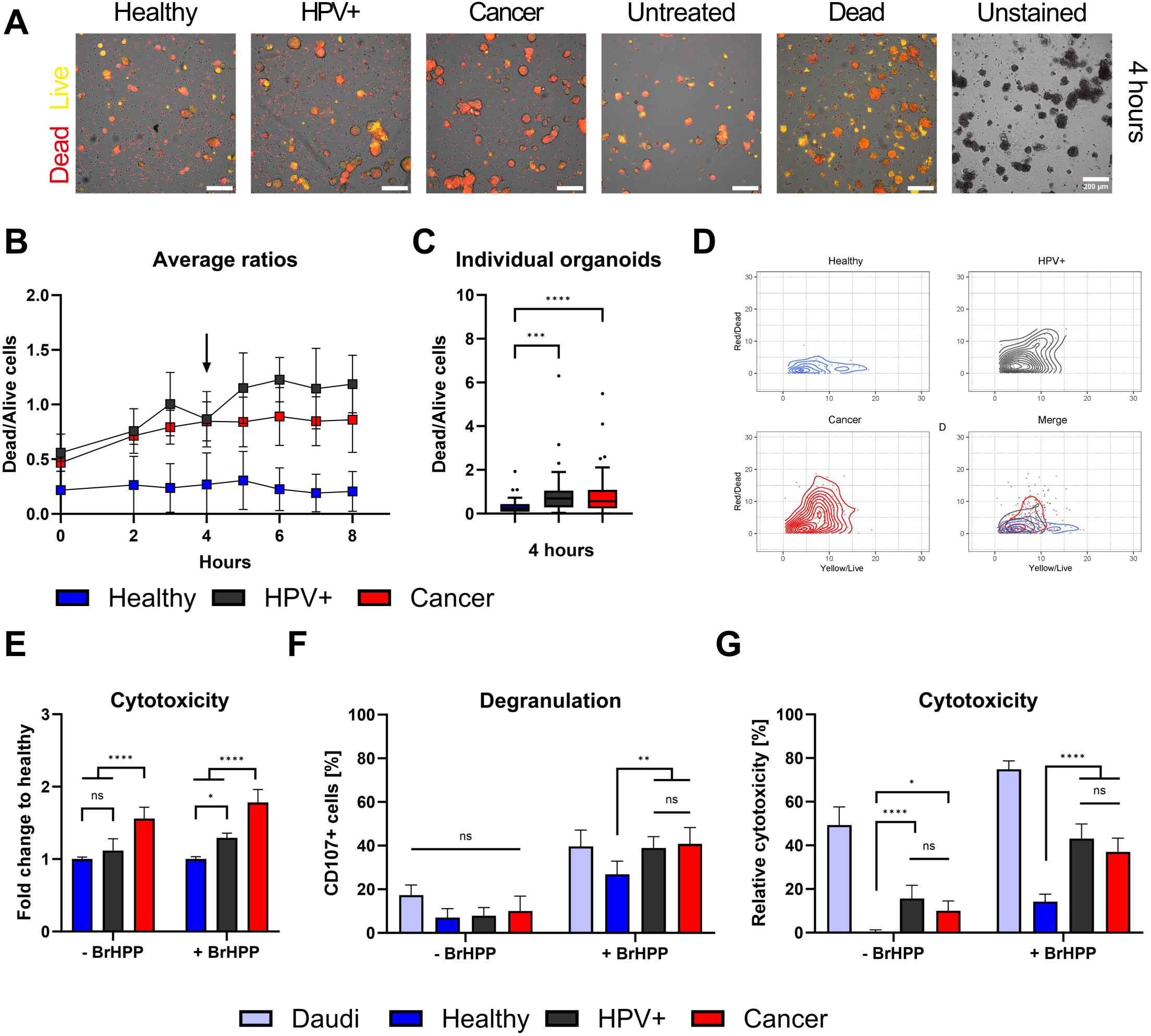
γδ T cells specifically target HPV-positive and cervical cancer organoids. A) Representative fluorescent images of ectocervical organoids co-incubated with γδ T cells for four hours. Cells were stained with NucRed Dead 647 Ready Probe Reagent. Yellow marks viable cells, red marks dead cells. Scale bars represent 200 µm. The experiment was performed at least twice with similar results. B) Quantification of a representative live cell imaging experiment in A) over time. Mean (± SD) of the ratio of dead/live (red/yellow) cell staining from ≥ 2 wells is shown. C) Ratio of dead/live (red/yellow) cell staining of individual organoids after four hours. Organoids with a ratio > 10 were excluded from the graph for visual purposes. Data shows ≥ 40 organoids per condition using Tukey whiskers. Statistical significance between conditions was determined using a Kruskal-Wallis test with Dunn’s correction for multiple testing. *** p < 0.001 **** p < 0.0001 D) Scatter plots of dead/live (red/yellow) cell staining in individual organoids show the increasing shift of yellow to red staining of HPV+ and cancer organoids. Merge of all conditions is seen at the lower right. Control conditions are depicted in Supplementary Figure 3. E) Quantification of cytotoxicity measured by MultiTox-Glo Multiplex Cytotoxicity Assay after four hours of co-incubation of organoids with γδ T cells in the absence or presence of 300 nM BrHPP as indicated. Data was normalized to untreated or BrHPP-treated controls and the ratio of dead/live cells was calculated; fold-change to healthy control was assessed. Data is presented as mean (± 95% CI) of 12-20 replicates per condition from two independent experiments. F) Quantification of degranulation of γδ T cells by CD107a staining with and without additional activation of γδ T cells by BrHPP using flow cytometry. The B lymphoma cell line Daudi was included as a positive control. Data shows mean (± 95% CI) of four independent experiments run in triplicates. G) Quantification of cytotoxicity induced by γδ T cells by propidium iodide (PI) staining with and without activation of γδ T cells by BrHPP using flow cytometry. Data shows mean (± 95% CI) of ≥ three independent experiments run in triplicates. Statistical significance in E-G was determined using a Two-Way ANOVA with Tukey’s correction for multiple testing. ns not significant * p < 0.05 ** p < 0.01 *** p < 0.001 **** p < 0.0001.

To confirm the differences in γδ T-mediated cytotoxicity induced by γδ T cells against healthy and HPV+ and cancer-derived cervical organoids, we determined degranulation/Lamp-1 mobilization of γδ T cells by flow cytometry. As positive control we included the B lymphoblast cancer cell line Daudi. As expected, we could show that addition of BrHPP significantly enhanced degranulation in all conditions (Fig. 2F, right). However, addition of BrHPP also resulted in a significant increase in degranulation of HPV+ and cancer organoids in comparison with healthy controls (Fig. 2F). Confirming our previous results using an alternative cytotoxicity read-out, we demonstrated that unstimulated γδ T cells already exhibit significantly increased cytotoxic activity against cancer and HPV+ cells in comparison to healthy controls as measured by propidium iodide (PI) staining to detect dead cells (Fig. 2G). Again, the addition of BrHPP increased the overall response and significantly enhanced the killing of HPV+ and cancer organoids as compared to healthy cells (Fig. 2G).

### Transcriptomic and protein analysis reveals differences in molecules relevant for γδ T cell activation

We have observed significant differences in γδ T cell-mediated cytotoxicity against distinct organoid lines. To identify putative underlying factors facilitating increased cytotoxicity on cancer and HPV+ organoids, explorative bulk RNA-sequencing (RNAseq) was conducted. Gene expression analysis revealed differences between conditions and also between patient isolates (Fig. 3A, Supplementary Data S1). Both cancer lines exhibited comparable expression of genes that were distinguishable from the other conditions, as shown by the first principal component (Fig. 3A, Supplementary Fig. 4). Isolates transformed with HPV oncogenes E6E7 (Supplementary Fig. 4, 5) grouped as intermediates between healthy and cancer organoids (Fig. 3A). Differential gene expression analysis showed a strong upregulation of cell proliferation associated genes in the HPV transformed isolates (Fig. 3B, Supplementary Data S1). In contrast, cancer organoids showed an upregulation of pathways involved in DNA-damage, cell cycle checkpoint, and DNA-binding/modification pathways and downregulation of pathways involved in differentiation of squamous epithelia (Fig. 3B, Supplementary Fig. 5; Supplementary Data S1). No significant elevation of antiviral/IFN-associated pathways was detected.

**Figure 3:**
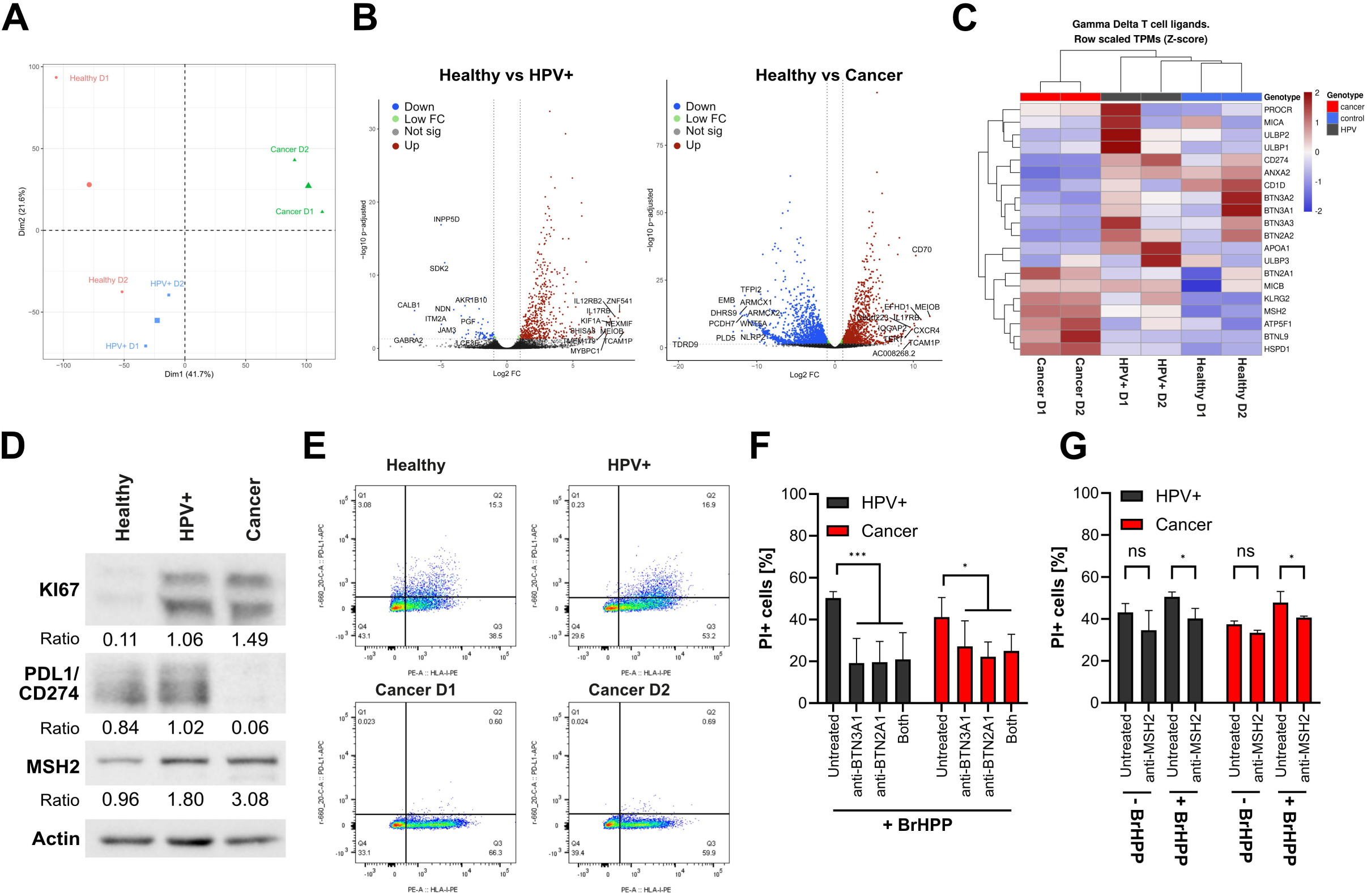
γδ T cell mediated killing is partially mediated by BTN family members and MSH2. A) PCA analysis of bulk RNA sequencing of the organoid lines. B) Volcano plots of differential gene expression analysis, showing mean Log2 fold-change vs −log10 adjusted p values. HPV16 integration alters the organoid transcriptome. RNA sequencing was performed for 2 independent patient organoid isolates per condition. Downregulated genes are marked in blue, differentially regulated genes with higher p values in green, not significantly regulated genes in grey, and upregulated genes in red. Top10 up- and downregulated genes are given. C) Heatmap of the gene expression of γδ T cell ligands highlighting the downregulation of BTN family members and the upregulation of DNA damage associated molecules. D) Confirmation of RNAseq results by Western blotting highlighting the differential expression of selected proteins in cancer organoids. E) Dot plot of PDL1/CD274 (Y axis) and HLA class I (X axis) expression analysed by flow cytometry on organoids as indicated, confirming protein expression results in D). F) Quantification of cytotoxicity in organoid lines by propidium iodide staining with and without blocking antibodies for BTN2A1/3A1 by BrHPP activated γδ T cells using flow cytometry. Data shows mean (± 95% CI) of three independent experiments. G) Representative quantification of a flow cytometry experiment using anti-MSH2 antibody (2-4 replicates per condition). The percentage of propidium iodide-positive cells is shown. Statistical significance in F and G was determined using a Two-Way ANOVA with Tukey’s correction for multiple testing. ns not significant * p < 0.05 *** p < 0.001

Having gained insights into the differential expression of transcripts between conditions we further explored the possible underlying mechanisms of enhanced killing of HPV+ and cancer organoids. For this, we investigated the expression of putative γδ T cell ligands and molecule involved in γδ T-cell activation (Fig. 3C) as summarized in Kabelitz, et al. ^11^. As expected, we observed pronounced differences in expression patterns between the conditions. Upon transformation with E6E7 gene expression patterns of healthy organoids shifted more torwards the cancer direction. Some molecules associated with γδ T cell activation such as the *Butyrophilin (BTN)*-family, *UL16 Binding Protein (ULBP)*-family, *MHC Class I Polypeptide-Related Sequence A (MICA), Apolipoprotein A1 (APOA1),* and *protein C receptor (PROCR)* were not elevated in cancer organoids. In contrast, others such as *MICB*, and *ATP Synthase Peripheral Stalk-Membrane Subunit B (ATP5F1/BP)* were expressed in a higher amount (Fig. 3C). *CD1d* and *Annexin A2 (ANXA2)* were downregulated compared to controls (Fig. 3C).

In addition to the well characterized genes involved in γδ T cell activation, we also investigated the regulation of other genes such as members of the *human MutS homologue (MSH)*-family, *heat shock protein family D (Hsp60) member 1 (HSPD1)* and *Killer Cell Lectin Like Receptor G2 (KLRG2)*. These genes have been described to play a role in DNA mismatch repair or are involved in stress signalling of cells ^24,25^. Interestingly, ectopically expressed *MSH2* has been previously identified as a target for human γδ T cells ^11,24,26,27^. We found these genes to be upregulated in HPV+ and cancer organoids (Fig. 3C). Another highly upregulated gene was found to be *Butyrophilin-Like Protein 9 (BTNL9)* which has been implicated to be of prognostic significance in various cancer types ^28,29^. Its potential involvement in the regulation of γδ T-cell mediated killing of tumour cells is unknown but deserves further investigation in light of our current findings with HPV-transformed and cancer organoids. Additionally, *programmed cell death ligand 1 (PD-L1/CD274)* was significantly downregulated in cancer organoids. The PD-L1 receptor PD1 conveys a negative signal on T cells. ^30^. PD-L1 downregulation might therefore support the enhanced killing of cancer cells by γδ T cells.

To confirm whether the regulation of transcripts could be translated into protein expression, we analysed the protein expression of selected differentially expressed genes by western blotting (Fig. 3D). We confirmed that KI67 was elevated in cancer-derived organoids, consistent with earlier reports ^31^ and as was observed in immunofluorescence (Fig. 1D). Additionally, expression of MSH2 was upregulated in HPV+ and cancer organoids and PD-L1 was downregulated in cancer organoids in comparison with healthy controls, pointing to possible functional differences in γδ T cell-mediated killing. As PD-L1 needs to be expressed on the cell surface to fulfil its function, we also conducted flow cytometry to confirm our western blotting and RNA-sequencing results, which, indeed, confirmed decreased surface expression of PD-L1 in cancer organoid isolates (Fig. 3E).

Next, we addressed the functional relevance of some of the above molecules for γδ T cell mediated killing of HPV+ and cancer organoids. To this end, we co-incubated organoids with expanded γδ T cells with BrHPP in the absence or presence of blocking antibodies against BTN2A1 and BTN3A1 (Fig. 3F). We found that blockade of BTN2A1 or/and BTN3A1 significantly reduced killing in all conditions as was in line with earlier reports ^32^. However, blocking in cancer organoids seemed to be slightly less affected than in HPV+ organoids (Fig. 3F). In view of the results shown in Fig. 3C/D, we also addressed the potential role of MSH2 which has been identified by Dai, et al. ^24^ as a γδ T-cell ligand when ectopically expressed on the surface of tumour cells. Blocking of MSH2 by antibodies led to an overall decrease of γδ T cell-mediated cytotoxicity, most notably against cancer organoids in the presence of BrHPP (Fig. 3G). Blocking could decrease γδ T cell-mediated cytotoxicity of BrHPP activated γδ T cells to the level of unstimulated γδ T cells (Fig. 3G). While the inhibitory effect of anti-MSH2 antibody blocking was much lower than the effect of anti-BTN2A1/3A1 antibodies, our results suggest that multiple ligand-receptor interactions are involved in the organoid-γδ T-cell interaction.

In summary, we have identified multiple possible target molecules and pathways that may contribute to enhanced killing of HPV-infected cervical tissue by γδ T cells. While the precise role of some molecules (including BTNL9) requires further investigation, the currently established organoid-γδ T-cell co-culture system should prove useful for testing additional modulators (e.g. small molecules, antibodies, etc.) with the ultimate goal of improving the efficacy of γδ T-cell immunotherapy for cervical cancer.

## Discussion

The association between HPV infection and the onset and the development of cancer has been extensively studied since almost 50 years ^33^. However, there has been a lack of suitable models that accurately represent and maintain the characteristics of cervical tissue. Previous studies have relied on immortalized cell lines or tissue slices ^18,20^. Here, we present a model to study the interaction of tumour-reactive T lymphocyte populations with healthy, matched HPV16 E6E7 transformed, and cancer-derived primary ectocervical tissues. To achieve this, we utilized an earlier established human ectocervical organoid model including paired HPV16 E6E7 transformed organoids ^5,6^. We detected infection-specific differences in proliferation patterns of healthy and HPV-infected organoids by staining with KI67 antibody. Healthy controls exhibited proliferating cells only in the parabasal regions of organoids, cancer organoids showed a wider distribution of proliferation while HPV+ showed an intermediate phenotype. Similar was observed in HPV+ tissue and cancer biopsies confirming the preservation of tissue origin characteristics ^31,34^.

γδ T cells play a distinct role in the immune surveillance against virally infected (including HPV) cells and tumour cells ^3,8,11,35^. They are believed to be important in the cervical defence against HPV infection ^3,36^. To establish an *in vitro* model for in-depth analysis, we co-incubated organoid-derived cells with *in vitro* expanded γδ T cells isolated from healthy donors. In view of the HLA independence of γδ T cells, the use of allogeneic γδ T cells from healthy donors is a valuable approach also from a translational perspective. In fact, the adoptive transfer of allogeneic γδ T cells from healthy donors has already been applied in phase I studies with no adverse reactions ^10^. It was thus possible to compare responses of γδ T cells to healthy, HPV-transformed as well as tumour organoids. Our results demonstrated elevated responses of unstimulated and activated γδ T cells to HPV-infected cells in comparison to healthy controls, independently of the type of assay used. Live cell imaging, protease-dependent cytotoxicity assays, and flow cytometry-based cytotoxicity and degranulation assays demonstrated elevated responses to HPV+ and/or cancer organoids.

To unravel possible mechanisms behind the responses of γδ T cells to HPV16-infected organoids, we conducted bulk RNA sequencing. The RNAseq revealed differences between HPV infected vs healthy controls. Importantly, HPV-transformed organoids displayed an intermediate state between healthy and cancer organoids with replicating some, but not all patterns of cancer tissue. Consistent with earlier studies we detected evidence for HPV16-induced DNA damage ^5,37^. Interestingly, HPV-induced antiviral responses including the type I interferon pathway were negligible in the pathway analysis. Dekkers, et al. ^23^ have earlier reported a high association between cancer cell killing and the preservation of tumour-specific inflammatory features in breast cancer organoids. In their study, sensitivity to cancer metabolome-sensing engineered T cells (TEGs) was highly associated with expression of antiviral genes such as *MX1*, *IFIT1*, *OASL* and *XAF1* ^23^.

Our pathway analysis demonstrated some overlap in DNA-modification pathways and γδ T cell ligands or associated molecules between cancer and HPV+ conditions. We reproduced earlier reports ^38,39^ of altered transcription and expression of genes such as *TTK protein kinase (TTK)*, *maternal embryonic leucine zipper kinase (MELK)*, *forkhead box M1 (FOXM1)*, *mismatch repair enzyme as mutS homolog 2 (MSH2)*, *matrix metalloproteases (MMPs)* and *programmed cell death ligand 1 (PD-L1/CD274)*. While up-regulation of *PD-L1* is typically associated with poorer outcomes in chemotherapy treatment ^39^, it was almost absent in our cancer isolates. While exhaustion and the role of PD-1, TIM3, LAG-3 and other checkpoint molecules has been less studied in γδ T cells compared to conventional αβ T cells ^40^, the low to absent PD-L1 expression on cancer organoids may favour persistent effector activity of γδ T cells. Interestingly, the expression of BTN-family members including BTN2A1/3A1, crucial in pAg-mediated γδ T cell activation was largely down-regulated in the cancer isolates. However, our blocking studies using inhibitory antibodies against BTN2A1 and BTN3A1 clearly indicated the significance of these BTN molecules in triggering the cytotoxic activity of γδ T cells against HPV+ and cancer organoids., in line with previous reports ^32^. While not directly measured in our study, it can be assumed that HPV+ and cancer organoids produce increased amounts of endogenous pAg isopentenyl pyrophosphate (IPP), leading to BTN-dependent γδ T-cell activation and subsequent tumour cell killing ^8,11,41^. Additionally, transcripts of, *BTNL9*, and cellular stress molecules *HSPD1, KLRG2,* and DNA damage associated molecule *MSH2* were upregulated in cancer organoids. While the roles of *HSPD1* and *KLRG2* play a role in regulating γδ T cell-mediated cytotoxicity in cancer have been described ^42,43^, there is limited information about the involvement of *BTNL9* in γδ T cell-mediated killing of cancer cells. Given our results, further investigation is warranted. Moreover, the upregulation of *MSH2* may be functionally relevant. MSH2 ectopically expressed on the surface of tumour cells has been identified as a ligand for the human Vγ9Vδ2 TCR ^24,26,27^ Our study demonstrated that the killing of HPV+ and cancer organoid cells by BrHPP-activated γδ T cells was reduced in the presence of anti-MSH2 antibody, supporting the hypothesis that the increased levels of MSH2 contribute to sensitizing of HPV+ and cancer organoid cells to cytotoxicity mediated by γδ T cells. Taken together, our findings suggest that BTN2A1/3A1 and MSH2 are involved in triggering the cytotoxic activity in γδ T cells towards HPV+ and cancer organoids.

In summary, we have established a suitable model to study cellular immune responses to healthy, HPV-transformed and cancerous ectocervical tissue using organoids in combination with short-term expanded γδ T cells. Co-incubation of organoid-derived cells with γδ T cells resulted in enhanced killing of HPV-transformed and cancer organoids. We demonstrated the involvement of BTN-family members BTN2A1/3A1 in γδ T cell-mediated cytotoxicity and obtained evidence for the additional involvement of human MutS homologue (MSH)-family member MSH2. The presented *in vitro* model will be highly valuable for in-depth analysis of additional pathways and mechanisms of how to enhance γδ T-cell effector function with the ultimate goal of improving γδ T-cell immunotherapy of cervical carcinoma.

## Acknowledgements

We thank Saskia F. Erttmann for proofreading the manuscript and valuable discussions. We thank Daniela Wesch, Institute of Immunology, UKSH Kiel for her support in obtaining leukocyte concentrates from blood bank donors. We thank Wiebe Schaefer, Reinhild Geissen, and Anna Willms from SYNENTEC GmbH for the provision of the Cellavista 4 automated cell imager, the SYBOT X-1000, the CYTOMAT 2 C-LiN system, and technical assistance throughout the live cell experiments. We thank ImCheck Therapeutics for kindly providing antagonistic anti-BTN2A1/3A1 antibodies. Overview schematics were assembled in BioRender.

## Authors contributions

Experiments were conducted and analysed by JD and DH. SK, RKG and CC established ectocervical organoids and transformed them with HPV E6E7 oncogenes. ME analysed flow cytometry data. CP and MZ performed and supervised flow cytometry experiments. MM provided tissues. NH established protocols and supervised live cell imaging experiments. HB and AQO analysed RNAseq data. AQO analysed live imaging data. DH drafted the manuscript. CC, DK and TFM conceived the project and designed experiments. All authors critically interpreted the data and revised the manuscript for important intellectual content. All authors contributed to the article and approved the submitted version.

## Material and methods

### Isolation of cervical tissue and cervical organoid culture

Human cervical specimens were obtained from volunteers undergoing standard surgical procedures at the Department of Gynaecology, Charité University Hospital, and August-Viktoria Klinikum, Berlin (Ethics Approval EA1/059/15 from local authorities), informed consent was obtained from all donors. Samples were processed within 2-3 hours after removal.

Organoids were derived from tissue resection as previously described ^5,6,22^. In short, biopsies were washed and cells were isolated by mincing and enzymatic digestion with 0.5 mg/mL collagenase type II for 2.5 h at 37°C. Afterwards, cells were pelleted at 100 x g for 5 min (4°C) and then again digested with TrypLE Express Enzyme (Gibco, 12604013) for 15 min at 37°C. Cells were then expanded in collagen-coated T25 tissue culture flasks in cervical organoid medium (Advanced DMEM/F-12 (Gibco, 12634) supplemented with 10 mM HEPES (Capricorn, HEP-B), 1% GlutaMax (Gibco, 35050061), 1x NCS21 Neuronal Supplement (Capricorn, C21-H), 1x N2 (Capricorn, N2-K), 10 ng/mL human EGF (Peprotech, AF-100-15), 0.5 μg/mL hydrocortisone (Sigma, H0888), 100 ng/mL human noggin (Peprotech, 120-10C), 100 ng/mL human FGF-10 (Peprotech, 100-26-25), 1.25 mM N-acetyl-L-cysteine (Sigma, A9165), 2 μM TGF-β receptor kinase Inhibitor IV (A83-01, Cayman Chemicals, Cay9001799), 10 μM ROCK inhibitor (Y-27632, Cayman Chemicals, Cay10005583), 10 mM nicotinamide (Sigma, N0636), 10 μM forskolin (Sigma, F6886)) until reaching 70-80% confluency. After successful establishment, cells were harvested and ~20.000 cells were seeded into 50 µl Matrigel domes (Corning, 11543550).

Organoids were maintained by passaging every 1-2 weeks at a 1:2-1:5 ratio using enzymatic digestion with TrypLE followed by mechanical singularization of cells using a syringe with an 18G needle (BD medical, BDAM303129) as described earlier ^22^.

### Cell lines and 3T3-J2 irradiation

AGS0 cells (AGS cell line devoid of *parainfluenza virus 5* infection, kindly provided by Richard E. Randall, School of Biology, St. Andrews University) were maintained in RPMI 1640 medium (Capricorn, RPMI-HA) supplemented with 1 mM Sodium pyruvate (Capricorn, NPY-B), 1x Penicillin/Streptomycin and 10% fetal calf serum (Sigma, F7524). HeLa (ATCC CCL-2) and 3T3-J2 cells (kindly provided by Craig Meyers; Howard Green laboratory, Harvard University) were maintained in DMEM (Capricorn, DMEM-HPSTA) supplemented with 1x Penicillin/Streptomycin (Gibco, 15070063), 10 mM HEPES and 10% fetal calf serum (Sigma, F7524). The 3T3-J2 cells were irradiated with 30 Gy in a Gammacell 40 Exactor. After irradiation, 1×10^6^ irradiated 3T3-J2 were seeded per T25 Flask and incubated overnight until all cells attached to the surface.

### 2D cervical stem cell lines culture

Passage 0 epithelial cells were cultured in plastic cell culture vessels without an irradiated 3T3-J2 monolayer and seeded in a Collagen I (1:100, Fisher Scientific, A1064401) pre-coated T25 flask. After most epithelial cells attached to the surface, the medium was changed every 2-3 days until confluence reached 80-90%. The cells were then passaged at a ratio of 1:2. For the passage 0, the medium was removed and the cells were washed three times with PBS (Capricorn, PBS-1A) and dissociated with 1 ml TrypLE (for each T25 flask) incubating 10 min at 37°C. The cells were detached from the flask surface and the cell suspension was collected with 10 ml Advanced DMEM/F12. After centrifugation at 400g for 4 min at 4°C, the cell pellet was resuspended with cervical organoid medium and seeded on irradiated 3T3-J2 monolayers.

From passage 1, ectocervical epithelial cells were cultured with irradiated 3T3-J2 layer and the medium was refreshed every 2-3 days. When it reached 80%-90% confluency, the cells were enzymatically detached twice. The first time, 0.5 ml TrypLE was used for each T25 flask for 1 min to detach the 3T3-J2 feeder cells, and then the cervical epithelial cells were detached with another 1 ml TrypLE and incubated for 10 min.

### Whole mount immunofluorescence assays (IFAs) and microscopic analyses

Phase contrast and brightfield images were taken using an IX50 (Olympus) microscope with a 4x or 10x objective. Images were contrast adjusted and scale bars were added using FIJI (ImageJ) ^44^.

For fluorescent microscopy, organoids were processed as described before ^45^. In short, organoids were harvested and washed once in PBS, then incubated in cell-recovery solution (Corning, 47743-696) for 30 min, followed by an additional wash with PBS (Capricorn, PBS-1A), and fixed for 20 min in 4% para-formaldehyde (PFA; Carl Roth, P087). Cells were permeabilized with 0.1 M glycine (Carl Roth, 3908), 0.2% Triton-X100 (Carl Roth, 3051) in TRIS-buffered saline (TBS; 50 mM Tris-Cl (Carl Roth, 9090), pH 7.5, 150 mM NaCl (Carl Roth, 3957)) and incubated in blocking buffer consisting of 3% bovine serum albumin (Carl Roth, 3854), 1% normal goat serum (Capricorn, GOA-1A), 0.2% Triton-X-100, and 0.1% Tween-20 (Sigma, P9416) in TBS for 2 h at RT. Primary antibodies were added for incubation at 4°C overnight in blocking buffer and organoids were washed the next day four times with 0.2% Triton-X100, 0.1% Tween-20 in TBS. Secondary antibodies and 10 μg/ml Hoechst 33342 (Thermo Fisher, H1399) as nuclear stain were added, kept for 1 h at RT in the dark, followed by three additional washing steps with TBS. After a final wash with deionized water, orgnoids were mounted with Fluoromount-G (Southern Biotech, 0100-01) on glass slides. Images were taken using a cLSM 880 microscope (Zeiss), equipped with Plan-Apochromat 20x/0.8 M27 and C-Apochromat 40x/1.20 water M27 objectives and analysed with ZEN blue software (v3.5) and FIJI. Antibodies and dilutions are listed in Supplementary Table 1.

### Isolation of gDNA and real-time quantitative-Polymerase chain reaction (RT-qPCR)

For nucleic acid extraction, organoids were harvested, pelleted, and washed with ice-cold PBS. Matrigel was removed by incubation with cell-recovery solution for 30 min at 4 °C. Genomic DNA (gDNA) was isolated with the Quick-DNA Miniprep (Zymo, D3024) according to manufacturer’s instructions. Purity of DNA was assessed by A260/A280 absorbance ratio using an Infinite M200 Pro plate reader (Tecan). RT-qPCR was performed with a StepOnePlus (Agilent) using the Luna Universal qPCR Master Mix (NEB, M3003), and included initial enzyme activation for 3 minutes at 95°C, followed by 40 cycles of 20s at 95°C, 30s at 60°C and 20s at 72°C. A minimum of 30 ng gDNA was used per well. Melting curve analysis was performed to verify amplicon specificity. Relative expression was calculated using the ΔCT method, and HPV status was assessed by electrophoresis of the amplified products. For that, amplicons were separated on a 1.5% agarose gel containing SYBR Safe Nucleic Acid Gel Stain (Invitrogen, S33102) in 0.5x TRIS-Borat-EDTA (TBE)-buffer (40 mM Tris-Cl, 45 mM boric acid (Carl Roth, P010), 1 mM EDTA (Carl Roth, X986)). The following primers were used:

HPV16-for 5’-AGCTGTCATTTAATTGCTCATAACAGTA-3’ ^46^;

HPV16-rev 5’-TGTGTCCTGAAGAAAAGCAAAGAC-3’ ^46^;

HPV18-for 5’-CGAACCACAACGTCACACAAT-3’ ^46^;

HPV18-rev 5’-GCTTACTGCTGGGATGCACA-3’ ^46^;

GAPDH-for 5’-GGTATCGTGGAAGGACTCATGAC-3’ ^5^;

GAPDH-rev 5’-ATGCCAGTGAGCTTCCCGTTCAG-3’ ^5^.

Signal was recorded using a ChemiDoc MP Imaging System (Biorad).

### Vδ2Vγ9 T cell expansion

Vδ2Vγ9 T cells were expanded from peripheral blood mononuclear cells (PBMCs) isolated from leukocyte concentrates of healthy blood donors using a Ficoll-Hypaque (Biochrom, Biocoll L 6113/5) density gradient according to the manufacturer’s protocol. Leukocyte concentrates were provided by the Institute of Transfusion Medicine UKSH Campus Kiel and were used in an anonymized fashion and Ethics Approval D 434/22 was provided by local authorities. The cell pellet was resuspended in RPMI medium (Capricorn, RPMI-STA) containing 10 % FCS (Sigma, F7524). Before expansion, the initial Vδ2Vγ9 T population among CD3+ T cells was checked by staining PBMCs with anti-Vδ2-FITC and anti-CD3-APC. Samples with a Vδ2Vγ9 T cell ratio over 2% was deemed acceptable and the sample was further processed. Vδ2Vγ9 T cells were selectively activated by stimulating PBMCs with 5 μM zoledronic acid (Novartis) and 50 IU/mL IL-2 (Novartis) over the expansion period of 14 days essentially as described ^47^. IL-2 was supplemented every other day. Purity of Vδ2Vγ9 T-cell lines was checked with anti-Vδ9-FITC and anti-CD3-APC. Lines were used for experiments when Vδ2Vγ9T cells represented more than 90% of the total cell population. Antibodies and dilutions are listed in Supplementary Table 1.

### Vδ2Vγ9 T cell co-culture with cervical cells

The B-cell lymphoma Daudi cell line (ATCC, CCL-213) with known sensitivity to Vδ2Vγ9 T-cell killing ^48^ was included as positive control. All lines were washed with PBS and pelleted in 15 mL falcon tubes by centrifuging at 1400 rpm for 5 min. Cells were stained with PKH67 (Sigma, PKH67GL) green dye was prepared according to manufacturer protocol. After incubation at RT for 4 min in the dark, 7 mL RPMI medium containing 10 % FCS was added and incubated for further 1 min to stop the staining. Cells were used at a ratio of 1:10 (epithelial cells: T cells). Co-culture was performed for 4 hours with the respective surface blocking mAb, if appropriate, and thereafter cells were labelled washed in PBS, and taken up in sample buffer containing 0.2 μg/ml propidium iodide (PI) for 20 min at 4°C and analysed by flow cytometry. Data was then analysed with FlowJo v10. The absolute number of viable epithelial cells (FITC+PI−) was calculated with the formula derived from Sacchi, et al. ^49^:

Cytotoxicity = 100− [(% of living epithelial cells in co-culture with γδ T cells/% of living epithelial cells without γδ T) × 100]

### CD107a degranulation assay

50 × 10^3^ effector cells were cultured alone or together with 5 × 10^3^ tumour cells for 4 h at a 10:1 ratio in the presence of anti-CD107a-PE mAb (clone: H4A3; BD Biosciences) and monensin (3 μM; added after 1 h). Thereafter cells were washed twice in PBS and analyzed by flow cytometry. Where appropriate, the synthetic pAg bromohydrin pyrophosphate (BrHPP; Innate Pharma, IPH1101) was added at 300 nM to activate Vδ2Vγ9 T cells. Cells were immediately acquired on a BD FACSCanto II flow cytometer. Data was then analysed with FlowJo v10.

### Live cell imaging

Prior to co-culture, a flat-bottom 384 well-plate (black with clear bottom, Corning) was coated with Poly-2-hydroxyethyl methacrylate (poly-HEMA; Sigma, P3932) in 100% ethanol as described earlier ^50^. 30 µl poly-HEMA was added in each well and the plate was left to dry overnight under the culture hood. The plate was washed with PBS immediately before use.

The imaging protocol was adapted from previous studies ^23,51^. Organoid lines were cultured for around seven days and Vγ9Vδ2 T cells were expanded for 14 days as described above. Matrigel was removed by incubation with cell-recovery solution for 30 min at 4°C. A part of the suspension of organoids was taken and digested into single cells using TrypLE then the number of single cells were counted to estimate organoid equivalents. The organoids were washed with cold PBS and thereafter organoids were collected and stained with CellTrace CFSE Cell Proliferation Kit (Thermo, C34554) according to the supplier’s protocol. Then, Vγ9Vδ2 T cells were added again at a ratio 1:10 (epithelial cells: T cells). NucRed Dead 647 ReadyProbe Reagent (Thermo, R37113) was added to distinguish dead and live cells. Dead cell staining was controlled with 1 mg/ml of Puromycin (InVivoGen, ant-pr-1) as positive control. Measurements were performed using the CELLAVISTA 4 automated cell imager in combination with the SYBOT X-1000 with CYTOMAT 2 C-LiN system (all SYNENTEC). Wells were imaged every hour for a total of 8 hours and afterwards fluorescence data and images were extracted with YT-Software (SYNENTEC GmbH) using the Real Cytoplasm (2F) application. The settings were modified to detect all organoids in the green channel and subsequently, the average intensity of the yellow and red signal within each spheroid area was analysed. The following setting were used for different channels. Exciter: Blue (475/28) - Emissionfilter: Green Filter 530nm (530/43), Exciter: Green (529/24) - Emissionfilter: Amber Filter 580nm (607/70), Exciter: Red (632/22) - Emissionfilter: Far Red Filter 670nm (685/40). Scatter plots were assembled in R (v4.1.0) with ggplot2 (v3.4.0).

### Live/Dead cytotoxicity assay

1×10^4^ cervical organoid cells were seeded into Collagen I pre-coated 96-well plates without feeder cells. On the next day, cells were co-incubated with 1×10^5^ γδ T cells (EC:TC = 1:10) for four hours in Opti-MEM (Gibco, 11058021). To quantify dead/live cells, the MultiTox-Glo Multiplex Cytotoxicity Assay kit (Promega, G9272) was used according to manufacturer’s instructions. The values were normalized to the respective untreated control, and the ratio of dead/live cells was then normalized to the healthy organoids to obtain fold changes.

### SDS-PAGE and Immunoblotting

For SDS-PAGE analysis, organoids were harvested, pelleted, and washed with ice-cold PBS. Matrigel was removed by incubation with cell-recovery solution (Corning) for 1 h at 4°C. Cells were then lysed in RIPA-buffer (50 mM Tris-HCl pH 8.0, 1% w/v NP-40 (Sigma, 492018), 0.5% w/v sodium deoxycholate (Sigma, D6750), 0.1% w/v SDS (Carl Roth, 0183), 150 mM NaCl, and 5 mM EDTA). After assessing the protein content using the Pierce Rapid Gold BCA Protein Assay Kit (Thermo, A53225) samples were diluted with 6x Laemmli buffer (70% v/v Tris HCl pH 6.8, 10% w/v SDS, 30% (v/v) glycerol (Carl Roth, 3783) and 0.01% w/v Bromophenol blue (Sigma, B0126) containing 10% v/v b-mercaptoethanol (Carl Roth, 4227). Samples were boiled at 95°C for 5 minutes. 25 µg of protein per condition was blotted onto Amersham Protran nitrocellulose membranes (Fisher Scientific, 10600016) using standard techniques and transfer quality was validated by Poinceau S (Sigma, P3504) total protein staining. The membranes were blocked for 1 hour in 1x RotiBlock (Roth, A151) and incubated with primary antibodies overnight. After three washing steps membranes were incubated with respective peroxidase-conjugated secondary antibodies and signals were detected using chemiluminescence with a ChemiDoc MP Imaging System (Biorad). Antibodies and dilutions are listed in Supplementary Table 1.

### Analysis of Bulk RNA sequencing

Quality of raw reads was assessed by FastQC. Reads were mapped to the human reference genome (assembly GRCh37) using the splice aware aligner STAR (v2.7) ^52^ in two-pass mode. Quality GTF files downloaded from Gencode (v28) ^53^, were used to map reads into genes with FeatureCounts in order to obtain raw counts. From this matrix of raw counts, genes with no expression across samples were filtered out, and differential gene expression was performed using DESeq2 (v1.32.0) ^54^ in R ^55^. Volcano plots were generated using ggplot2 (v3.4.0). Genes were ranked based on their log2 fold change compared to healthy controls, and gene set enrichment analysis (GSEA) was performed using the fgsea implementation from the clusterProfiler R package (v4.0.5) ^56^. Gene sets from Hallmark (H), Curated Gene Sets (C2) and GO Biological Process (C5-BP), retrieved from MsigDB ^57^ using msigdbr (v7.5.1) were studied. The gene expression heatmaps of the TPMs were generated using the pheatmap package (v1.0.12). Principal Component Analysis (PCA) was performed on the TPM normalized counts and the plots were generated using factoextra (v1.0.7) on the result of the pca function in base R. All the analyses were performed in R (v4.1.0). RNA-seq DGE and GSEA analysis results are included in Supplementary Data S1.

### Statistical analysis and panel composition

Basic calculations were performed using MS EXCEL 2016 (Microsoft). Figures were plotted using Prism 9.5.1 (GraphPad) or R (v4.1.0). P values ≤ 0.05 were considered as statistically significant. Asterisks indicate statistical significance values as follows: * p < 0.05 ** p < 0.01, *** p < 0.001, **** p < 0.0001. The panel composition and annotations were created using Affinity Designer 1.10 (Serif).

