## Supplementary Data for "γδ T cell-mediated cytotoxicity against patient-derived healthy and cancer cervical organoids"

**Supplementary Figures:**

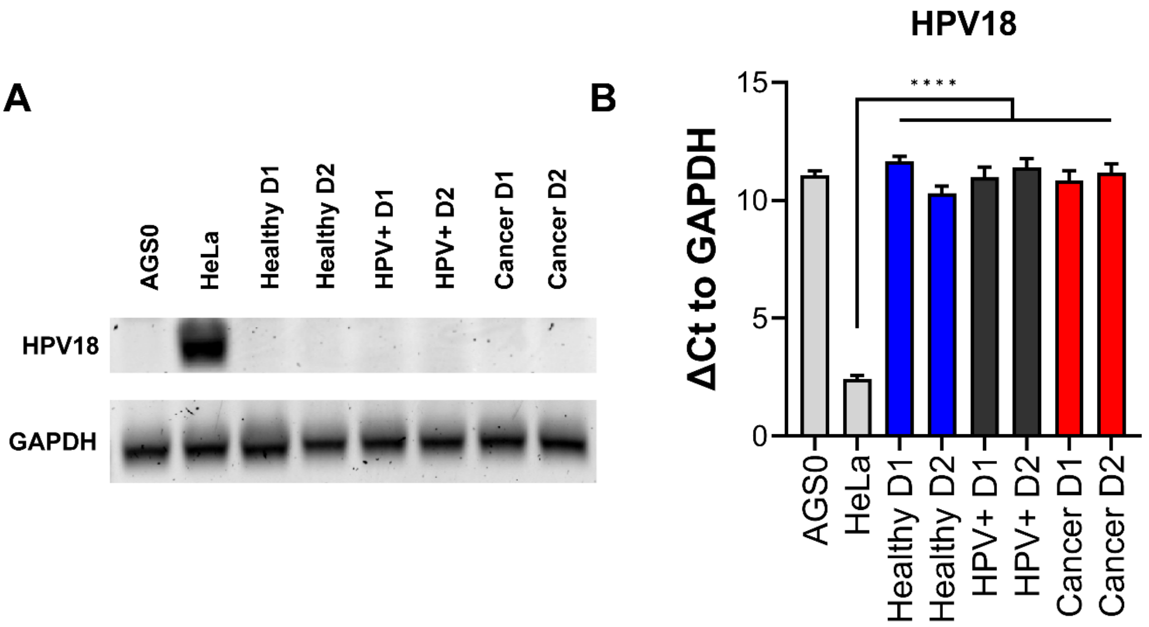


**Supplementary Figure 1: Summary of HPV18 status used patient-derived organoid cultures.** *Genomic DNA was amplified by RT-qPCR and ΔCt values were calculated with GAPDH as loading control. Same amplicons were run on agarose gels and HPV status was assessed. AGS0 and HeLa cells served as negative and positive control, respectively. HPV+ and cancer organoids show no presence of integrated HPV18. RT-qPCR experiments show mean (± 95% CI) of ≥ 3 technical replicas of two independent biological replicates. Statistical significance between conditions was determined using a Two-Way ANOVA with Tukey's correction for multiple testing. **** p < 0.0001*

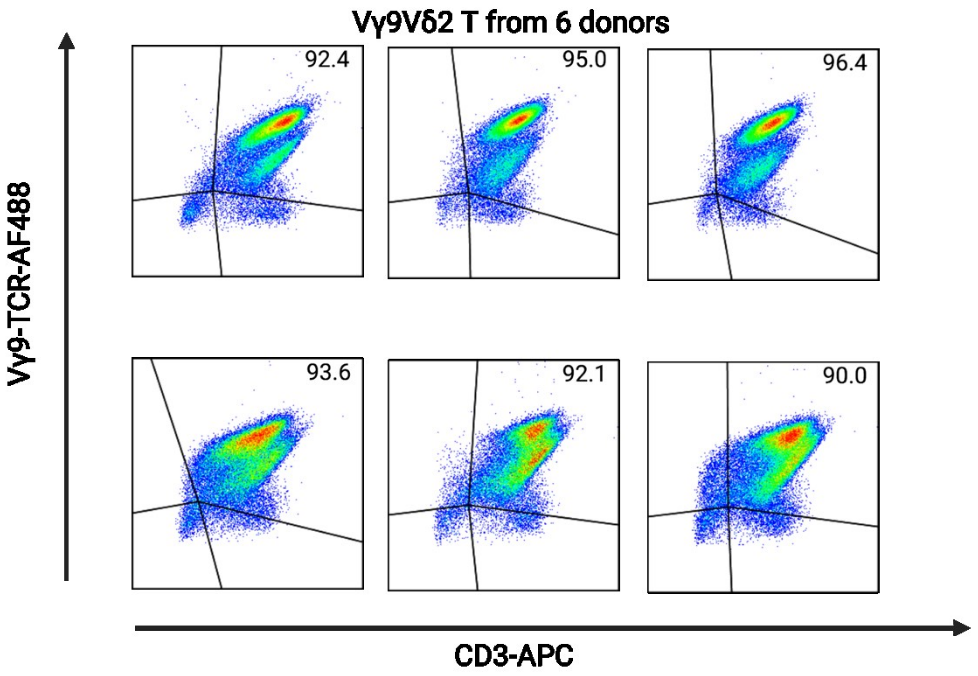


**Supplementary Figure 2: Exemplary purity measurements of γδ T cells by flow cytometry.** *Only cultures with a purity >90% were included.*

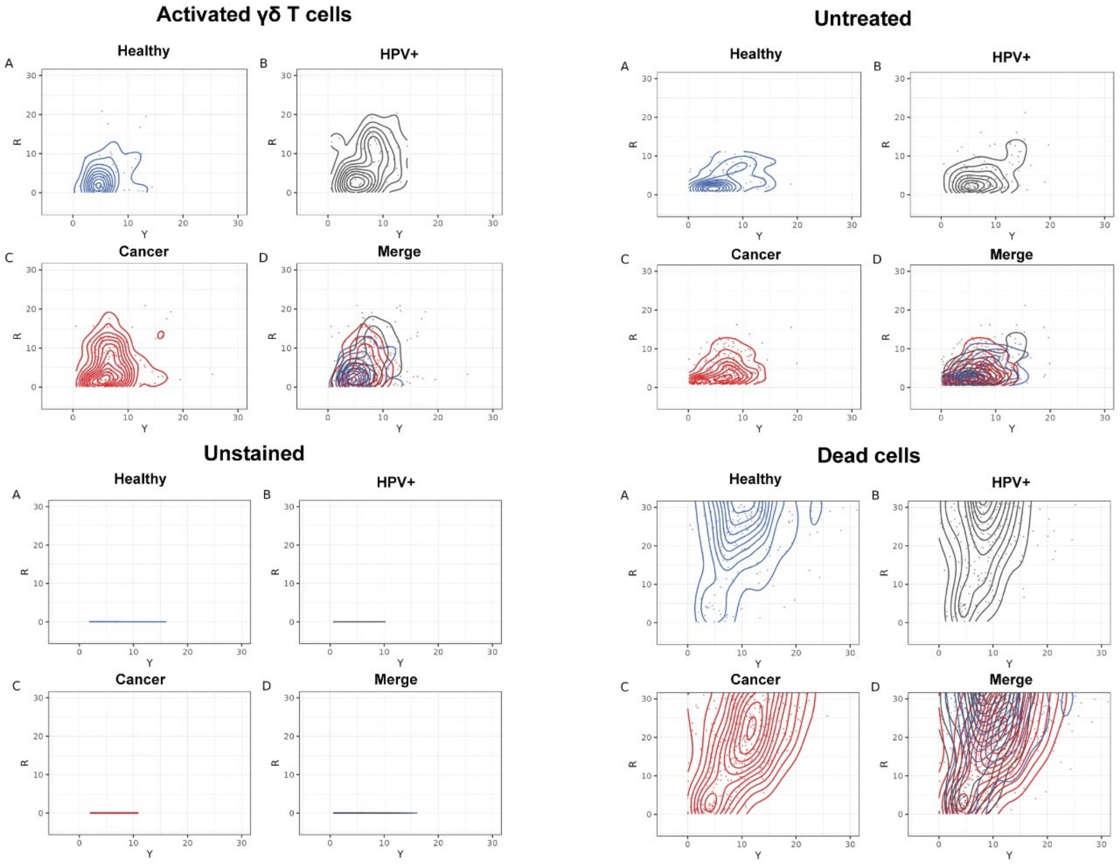


**Supplementary Figure 3: Scatter blots of live cell imaging depicting individual organoids after four hours in their respective conditions.** *Cells were stained with NucRed Live 647 Ready Probe Reagent. Yellow marks viable cells, red marks dead cells.*

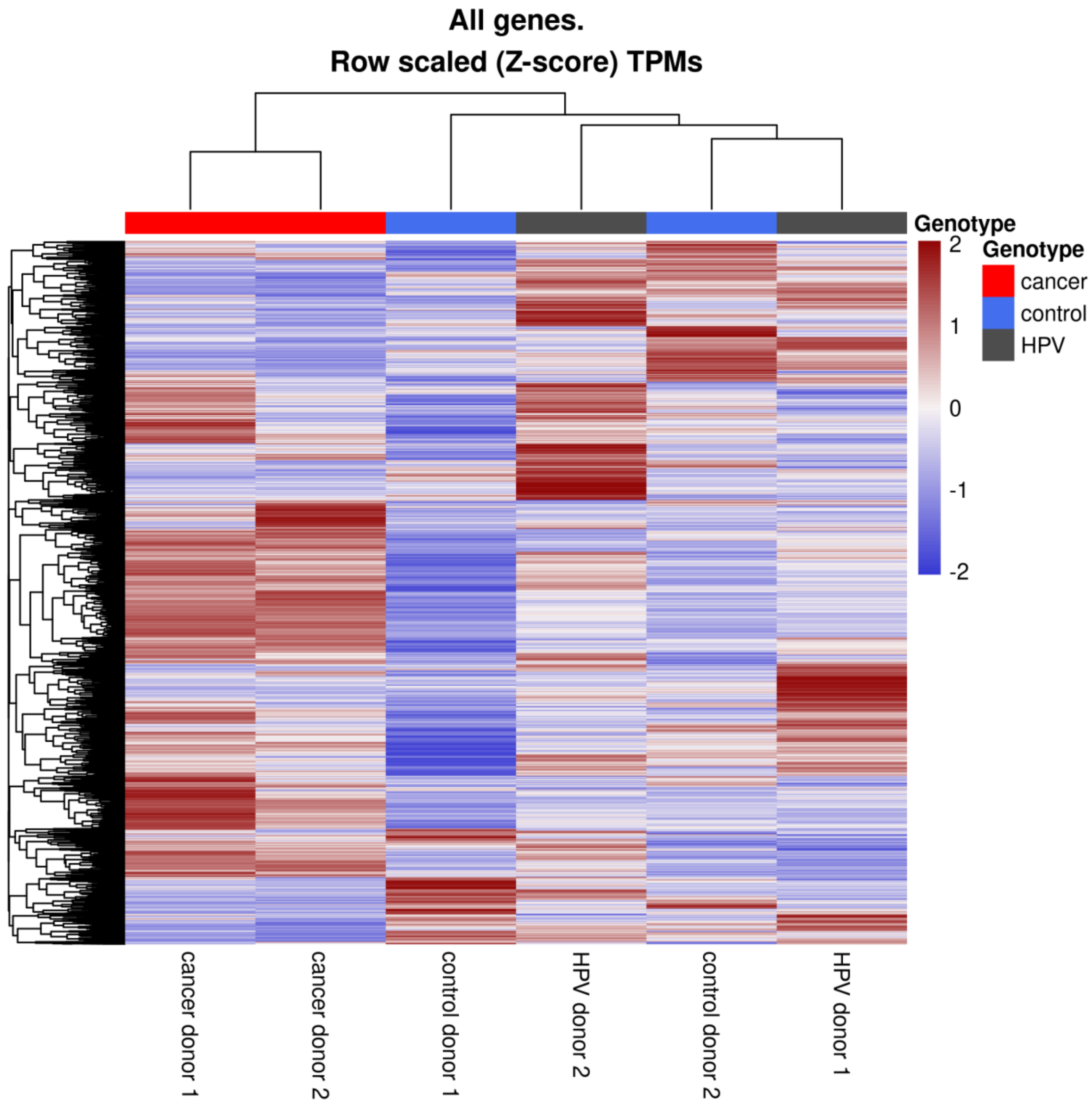


**Supplementary Figure 4: HPV16 integration alters organoid transcriptome**. *RNA sequencing was performed for 2 patient isolates per condition. Heatmaps of the z-score of normalized counts. Downregulated genes are marked in blue, and upregulated genes in red.*

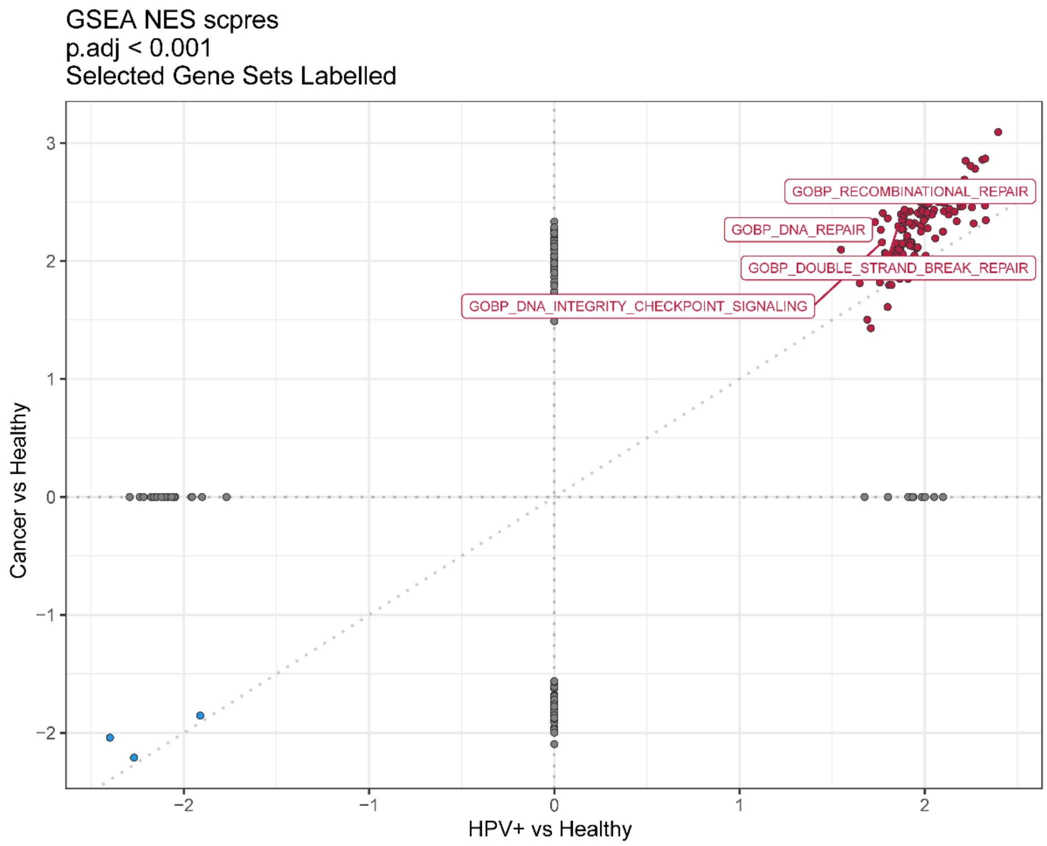


**Supplementary Figure 5: Significantly altered GOBP pathways as defined by GSEA analysis.** *Exemplary DNA altering pathways are given. Only Pathways below p adj < 0.001 are shown.*

**Supplementary table 1: Antibodies used in the study.**

| Target | Host | Vendor | Application | Dilution |
|---|---|---|---|---|
| **Primary antibodies** | | | | |
| KRT5-A488 | Rabbit | Abcam, ab193894 | IF | 1:500 |
| KI67 | Rabbit | Abcam, ab16667 | IF<br>WB | 1:500<br>1:1,000 |
| BTN2A1 | Mouse | ImCheck Therapeutics, mAb 7.48 | Blocking | 5 µg/ml |
| BTN3A1 | Mouse | ImCheck Therapeutics, mAb 103.2 | Blocking | 5 µg/ml |
| MSH2 | Rabbit | Cell Signaling, 2017S | WB<br>Blocking | 1:1,000<br>3 µg/ml |
| CD107a-PE | Mouse | BD Biosciences, Clone H4A3, 560948 | FC | 2.5 µg/ml |
| TCR Vδ2-FITC | Mouse | Beckman Coulter, Clone B6, 555738 | FC | 2 µg/ml |
| TCR Vδ9-FITC | Mouse | Conjugated in house, Janssen et al. (1991) | FC | 1:20 |
| CD3-APC | Mouse | BD Biosystems, Clone UCHT1, 561810 | FC | 2 µg/ml |
| HLA-ABC-PE | Mouse | BD Bioscience, 555553 | FC | 1:20 |
| PD-L1/CD274<br>PD-L1/CD274-APC | Rabbit<br>Mouse | Cell Signaling, 13684S<br>Invitrogen, 17-5983-42 | WB<br>FC | 1:1,000<br>1:20 |
| ACTB | Mouse | Sigma, A5441 | WB | 1:10,000 |
| **Secondary antibodies** | | | | |
| Anti-Rabbit A647 | Donkey | Jackson ImmunoResearch, 711-605-152 | IF | 1:100 |
| Anti-Rabbit HRP | Goat | Cell Signaling, 7074S | WB | 1:3,000 |
| Anti-Mouse HRP | Horse | Cell Signaling, 7076S | WB | 1:3,000 |
